# Comprehensive analysis of the internal structure and firmness in American cranberry (*Vaccinium macrocarpon* Ait.) fruit

**DOI:** 10.1101/567958

**Authors:** Luis Diaz-Garcia, Lorraine Rodriguez-Bonilla, Matthew Phillips, Arnoldo Lopez-Hernandez, Edward Grygleski, Amaya Atucha, Juan Zalapa

## Abstract

Cranberry (*Vaccinium macrocarpon* Ait.) fruit quality traits encompass many properties. Among these, fruit firmness has become a quality standard due to the recent demand increase for sweetened and dried cranberries (SDC). Traditionally, this trait has been measured by the cranberry industry using compression tests; however, it is poorly understood how fruit firmness is influenced by other characteristics. In this study, we developed a high-throughput computer-vision method to measure the internal structure of cranberry fruit, which may in turn influence cranberry fruit firmness. We measured the internal structure of 16 cranberry cultivars measured over a 40-day period. Internal structure data paired with fruit firmness values at each evaluation period allowed us to explore the correlations between firmness and internal morphological characteristics.

## 1. Introduction

The American cranberry (*Vaccinium macrocarpon* Ait.) is a native plant species from North America that has historically been associated with holiday traditions and distinctively tart and sweet juices (Song and Hancock, 2011; Vorsa and Johnson-Cicalese, 2012). Cranberries, açaí berries, blueberries and other berries are considered “superfruits” due to the vast amount of healthy antioxidants and phytochemicals they contain (Vorsa et al., 2000; Vvedenskaya and Vorsa, 2004; Wang et al., 2017). North America is the world’s largest producer of cranberries, led by the United States, which produces 58% (436,691 tons) of the world’s total cranberry production (FAOSTAT, 2016), with 95-97% of the fruit destined for processing and only 3-5% used for fresh fruit consumption (USDA NASS, 2017). In the US, the cranberry processing industry generates more than 11,000 jobs, and cranberry production supports a value added of $3.5 billion in economic activity (Alston et al., 2014; Jesse et al., 1993). However, in the last decade the US cranberry industry has faced economic and production challenges reducing the profitability of growers and processors. An over-production of juice concentrate and the requirement of labeling products with added sugar have recently reduced fruit prices considerably (USDA-ERS, 2017, U.S. FDA, 2016). Nevertheless, the popularity of sweetened and dried cranberry (SDC) has increased resulting in a growing demand. One of the main constraints for increasing SDC production is the strict fruit quality requirements for this production line. Desirable characteristics for SDC production include evenly colored, large size, round shape, and fully fleshed fruit with the ability to maintain high firmness as well as intact internal structure during processing after long-term frozen storage (Lindsay Well-Hansen, 2018, personal communication). A 2017 survey conducted at grower meetings in Wisconsin, New Jersey and British Colombia found that 53% of the respondents identified fruit quality traits as the most needed in the cranberry industry. The same respondents also indicated that fruit firmness was the fruit quality trait they believed to be the most important for breeding efforts. Fruit firmness is important to increase the production of SDCs, which are critical for the economic viability of the cranberry industry (Gallardo et al. 2018).

Moreover, cranberry fruit firmness must be considered not only during the industrial processing of SDC products, but also during fruit ripening and the post-harvest processes. Cranberry vines produce non-climacteric crisp fruit during the fall season, which undergo a moderate softening as they ripen (Ozgen et al., 2002). Excessive softening during fruit ripening can cause post-harvest deterioration, transportability issues, limited storability, and shelf-life decrease (Hadfield and Bennett, 1988). Commonly, cranberry fruits are stored in crates for up to three months at temperatures ranging from 1.7 to 4.4°C (Gunes et al., 2002). These post-harvest conditions may cause softening by mechanical damage deterioration, fungal fruit rot, or physiological decay (Forney, 2010, 2003; Meli et al., 2010).

Cell wall structure, turgor, cuticle properties and biochemical constitution have all been found to affect fruit firmness (Saladié et al., 2007; Chaïb et al., 2010; Chapman et al., 2012). In cranberry, fruit firmness is hypothesized to be intrinsically correlated with internal structure due to locular variability in the fruit. This may make internal structure attributes an important factor in fruit firmness. Unlike external physical attributes of the fruit such as color, size and shape, the internal structure properties have represented a challenge to measure, and until now, no method was available to measure them (Figàs et al., 2015; Hurtado et al., 2013; Yoshioka and Fukino, 2009, Diaz-Garcia et al., 2018). Also, the variability of internal structure characteristics across different cranberry cultivars and the contribution of internal fruit structure parameters to fruit firmness and shelf life are poorly understood (Gorny et al., 2008; Kumar et al., 2012; Zude et al., 2006).

In this study, we developed a computer vision, semi-automated method to quantitatively measure several parameters related to the internal structure of the cranberry fruit. Additionally, to better understand if firmness properties in cranberry fruit are affected by internal structure characteristics, we obtained compression measurements during the ripening process and explored the possible correlations. Lastly, we studied the effects of frozen storage on fruit firmness. This research contributes a method to define key parameters useful for SDC production.

## 2. Materials and Methods

### 2.1. Measuring internal structure characteristics in cranberry fruit

Our phenotyping method presented below involves dissected cranberry fruit being made into stamps using regular ink and paper. Stamping cranberry fruit is relatively easy, and it has already been implemented in our lab as a standard procedure to examine internal fruit structure. We developed a computer vision-based method to quantitatively measure fruit stamps in a high-throughput fashion. This process starts by making an equatorial cut at the widest section of the berry that is then coated in black ink and stamped on a sheet of paper. This imprint shows all four locules allowing their measurement as well as the quantification of other attributes such as the total pericarp area and fruit size. Fruit stamps can be made on regular white printer paper (which can contain Quick Response codes [QR] to easily automate further processing), and then digitized in a conventional scanner (Figure 1A). Subsequently, a custom MATLAB code processes each scan as follows. For each scanned image, individual stamps are identified and labeled using the Hough transformation (Figure 1B). Then, a circular mask is applied so that only the stamp is exposed; this also reduces the noise caused by bumps or imperfections in the stamping process. The pixels corresponding to the pericarp and locules are differentially labeled (see color bar in Figure 1B) based on a custom threshold and then quantified. Later, each stamp is rotated one degree at a time using nearest-neighbor interpolation, and the row-wise sum is iteratively computed (Figure 1C). Using this strategy, the rows with larger amounts of pericarp are going to add fewer quantities (pixels) compared with the ones that contain locules. This rotation/sum process is iterated until 180 degrees is reached (after this point, the output becomes identical). The results are converted into a matrix, denominated here as R (or rotational matrix), with a size *n* x 180, where *n* = stamp diameter (in pixels). Finally, the column-wise sum is computed using the R matrix, and a threshold is applied to determine the outer pericarp diameter (OPD). This approach allows the estimation of an average OPD across all the radial points of the stamp, which can be extremely useful when non-symmetric fruit stamps are measured. Using the OPD estimate, the external portion of the pericarp is masked and the internal pericarp is also quantified. Later on, other important parameters such as total pericarp area-to-locule size ratio are calculated based on the primary traits discussed here.

**Figure 1.**
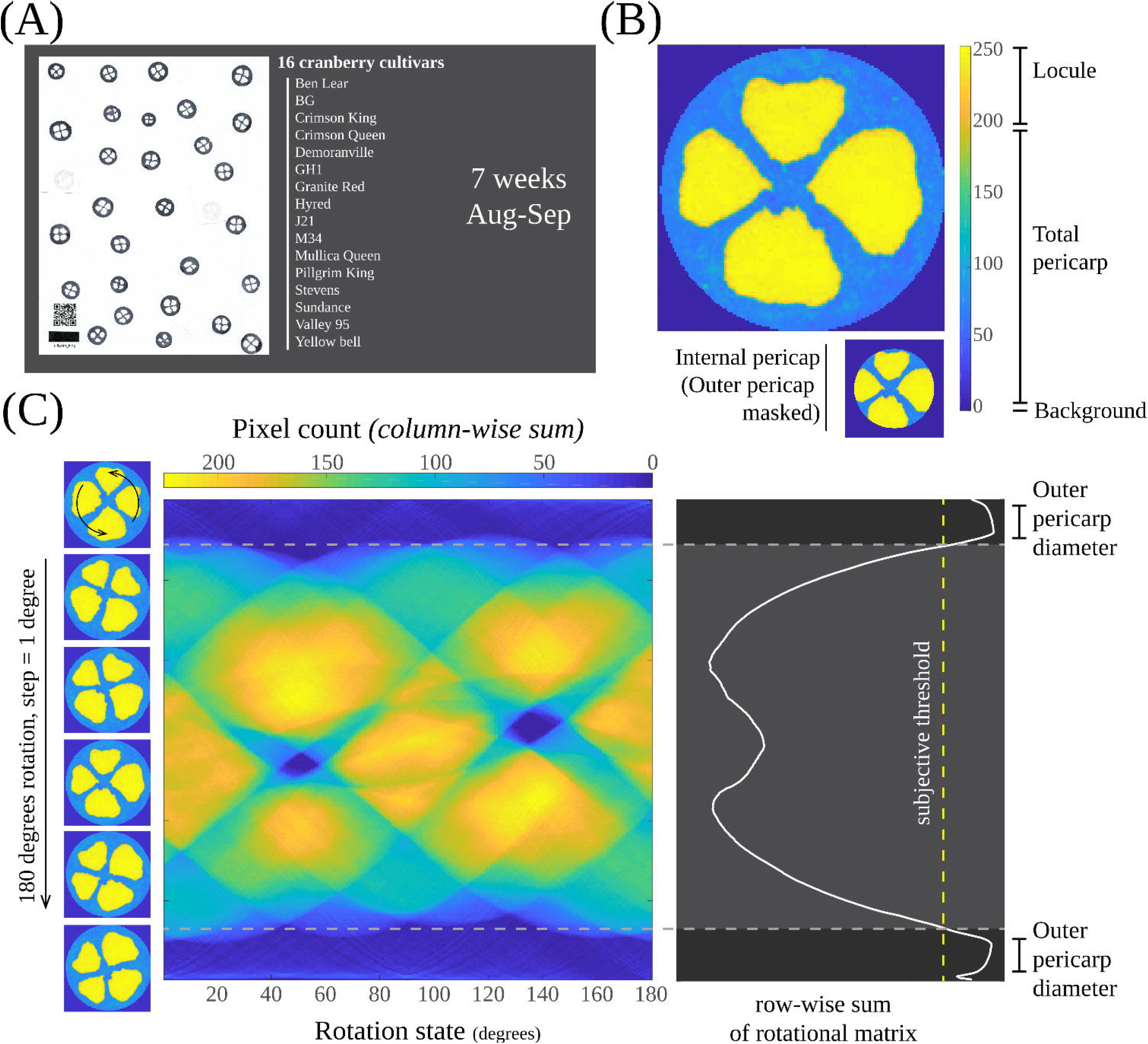
Fruit structure parameters measured through computer vision. (A) A set of fruit (N>25) from 16 cranberry cultivars were equatorially cut, and a single ink stamp per fruit was imprinted on paper. QR codes (for automating image processing) and size reference were included on each page. (B) Background, total pericarp area and locules were differentially flagged to estimate fruit structure parameters. (C) Outer pericarp diameter was obtained by computationally rotating each fruit stamp and analyzing the horizontal pixel distribution.

### 2.2. Phenotyping internal fruit structure in cranberry cultivars

Fruit were harvested during a 40-day period on dates separated by 6-10 day intervals and the internal fruit structure was phenotyped as described above. The measurements included the early and intermediate harvest seasons, and correspond to the following dates in 2017; July 27 (day 0), August 2 (day 6), August 9 (date 13), August 18 (date 22), August 23 (date 27), August 29 (date 33) and September 5 (date 40). The commercial cranberry cultivars used in this study were ‘Ben Lear’, ‘BG’, ‘Crimson King’, ‘Crimson Queen’, ‘Demorranville’, ‘Granite Red’, ‘GH1’, ‘HyRed’, ‘Mullica Queen’, ‘Stevens’, ‘Sundance’, and ‘Pilgrim King’. The experimental accessions tested were ‘J21’, ‘M34’, ‘Valley 95’, and ‘Yellow Bell’. ‘Stevens’ is the industry standard and most widely planted cultivar, accounting for over 40% of the total U.S. acreage (Cranberry Institute data, https://www.cranberryinstitute.org). ‘Mullica Queen’, ‘Sundance’, and ‘Crimson King’ are increasingly popular new varieties (Gallardo et al. 2018). ‘Pilgrim King’ has the largest cranberry fruit of any commercial cultivar while ‘Granite Red’ has the highest firmness and longest storage life among commercial cultivars (www.cranberryvine.com/cranberry-varieties). In addition, some of these varieties have traditionally been preferred by the industry for SDC (e.g., ‘Stevens’ [Lindsay Well-Hansen, 2018, personal communication]). During the evaluation process, we harvested a random sample of fruit from cranberry commercial beds planted in two different farms in Central Wisconsin, USA. At least 25 fruits per cultivar were used for stamps and 20 for firmness analysis (method described below). We also recorded the weight of 40 fruits per cultivar for fruit density estimation.

### 2.3. Firmness analysis

At each time point, we measured maximum compression force and maximum compression distance of 20 fruits per cultivar. These measurements were made on a Texture Analyzer (*TA.XTPlus Connect*, Textural Technologies, Hamilton, MA, USA) using 1mm·s^-1^ test speed, and “Strain” as target mode (30%). For a given compression profile (curve with force in the y axis and distance in the x axis), the traits maximum compression force and maximum compression distance corresponded to the maximum value reached by the curve, and the distance in which that maximum force was reached, respectively. In addition, for the last two harvest dates (33 and 40), 20 fruits per cultivar were stored at −25°C for 60 days and then thawed at room temperature, simulating the process used for SDC production. Using the same compression measurement protocol thawed fruit were also tested.

### 2.4. Statistical analysis

The overall fruit structure and firmness changes were analyzed in R (R Core Team, 2013), using all cultivars as repetitions, and by fitting linear quadratic models of the form *y~x+x2*, where *y* corresponded to the observed trait, and *x* to the time points. Subsequently, quadratic models of the same form were fitted for each cultivar and the model coefficients (*beta1* and *beta2*) were recorded for further comparisons. Additionally, using the predicted values (based on the previous linear models) during a 45-days period, a principal component analysis (PCA) was performed with the R function *prcomp*, and the three first components were recorded. Finally, a PCA was performed using all time points together, and the cultivars were visually analyzed by plotting the PC1 and PC2.

## 3. Results

### 3.1. High-throughput phenotyping of fruit stamps and internal structure variation across time

Using a high-throughput imaging method, we measured more than 3000 individual ink stamps representing 16 cranberry cultivars over a 40-days period (7 time points total). For each cranberry fruit stamp, our method allowed us to estimate the following internal fruit structure parameters (Table 1): fruit size (FS), total pericarp area (TPA), locule size (LCS), outer pericarp diameter (OPD), internal pericarp area (IPA), and total pericarp area-to-locule size ratio (TPA/LCS). To better characterize if internal structure affects the firmness properties of cranberry fruit, we measured compression force and distance at each measurement time point for all 16 cultivars. Also, we recorded fruit weight, which allowed us to estimate fruit density (using FS as a parameter to calculate fruit volume, assuming a spherical shape). Because of the destructive nature of these approaches, different sets of fruit were used for each analysis (e.g., stamps, firmness, and weight) and their means were used for comparison purposes.

**Table 1.** Properties of cranberry fruit measured in this study.

| Parameter | Description | Units | Formula | Abbreviation |
| --- | --- | --- | --- | --- |
| *Stamp-based descriptors* |  |  |  |  |
| Fruit size | Stamp major axis length (fruit diameter) | cm | - | FS |
| Total pericarp area | 2D pixel count corresponding to fruit pericarp (fruit flesh) | pixel count | - | TPA |
| Locule size | 2D pixel count corresponding to locule size (air pockets) | pixel count | - | LCS |
| Outer pericarp diameter | Diameter of external pericarp | cm | - | OPD |
| Internal pericarp area | 2D pixel count of internal pericarp | pixel count | - | IPA |
| Total pericarp area-to-locule size | Total pericarp compared with locule size | ratio | TPA/LCS | TPA/LCS |
| *Weight descriptors* |  |  |  |  |
| Fruit weight | Fruit weight (based on the weight of 40 fruits) | g | - | FW |
| Fruit density | Fruit density (based on stamp dimensions) | g/cm<sup>3</sup> | FW/FS | FD |
| *Firmness-based descriptors* |  |  |  |  |
| Maximum compression force | Force needed to compress fruit to 70% its original height | g | - | CF |
| Maximum compression distance | Change in fruit height after being compressed to 70% its original height | mm | - | CD |

Fitting linear models using all cultivars as repetitions allowed us to understand the overall dynamics of fruit structure development during the maturation process (Figure 2A). Although differences among cultivars made the overall distribution disperse, most of the terms in the fitted quadratic models were shown to be significant at *p*=0.05 (except for *beta2* in OPD). Predicted values from each quadratic model showed that fruit size ranged from 1.20 to 1.47cm (+23%), total pericarp area from 7653.38 to 12001.34px (+57%), locule size from 3702.99 to 5071.88px (+37%), total pericarp area-to-locule size from 2.21 to 2.48 (+12%), outer pericarp diameter from 0.16 to 0.22cm (+34%), internal pericarp area from 1496.07 to 2116.51px (+41%), compression force from 4556.36 to 6162.30g (+35%), compression distance from 4.79 to 6.03mm (+26%), fruit weight from 0.74 to 1.64g (+122%), and fruit density from 0.23 to 0.26g/cm^3^ (+16%).

**Figure 2.**
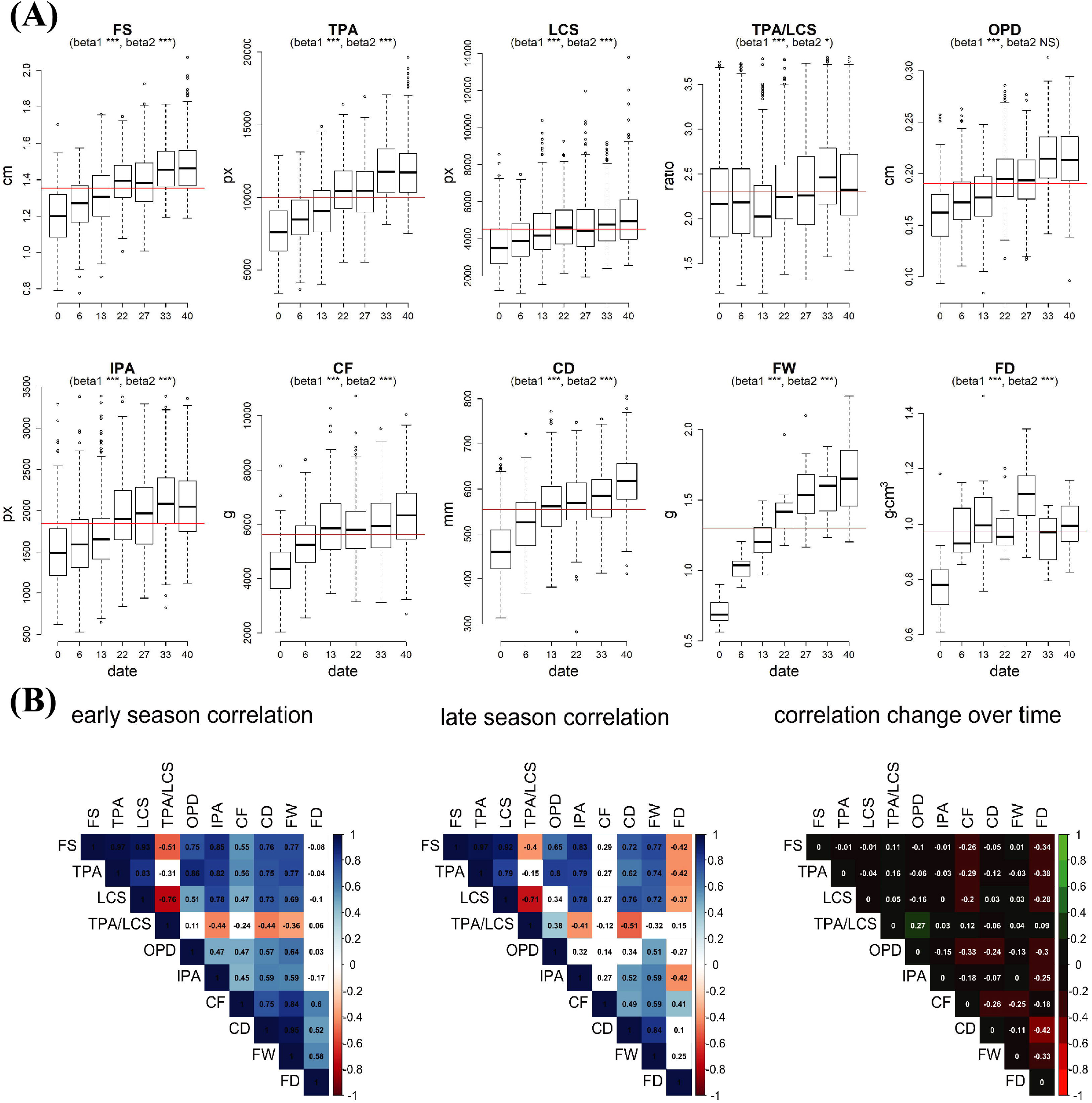
Variation in fruit internal structure and firmness. (A) Changes in fruit size (FS), total pericarp area (TPA), locular cavity size (LCS), total pericarp area-to-locular cavity size (TPA/LCS), outer pericarp diameter (OPD), internal pericarp area (IPA), compression force (CF), compression distance (CD), fruit weight (FW), and fruit density (FD), during the ripening process of cranberry. Quadratic models were used for each phenotype all cultivars as repetitions. On each boxplot, red line indicates the mean across all time points; significance (*** = *p* <0.005, ** = *p*<0.01, * = *p*<0.05, NS = *p*>0.05) of *beta1* (*x*) and *beta2* (*x^2^*) terms are provided in the boxplot subtitle. (B) Pearson’s correlation between fruit structure and firmness attributes at dates 0 and 6 (early season, first panel), and 33 and 40 (late season, second panel). Non-significant correlations (at *p*=0.05) are colored white. Third panel shows the changes in correlation when comparing late season and early season correlations.

Many of the fruit structure and firmness descriptors showed strong correlation with each other (Figure 2B). Although the correlation between most of the descriptors was consistent throughout the ripening process, we found that some traits correlated differentially when comparing early (first two dates) and late (last two dates) season data. Fruit size consistently had a Pearson’s correlation greater than 0.90 with both total pericarp area and locule size. Similarly, compression force and fruit weight showed a moderate correlation (0.62-0.77) at both season times with fruit size, total pericarp area, and locule size. Locule size and total pericarp area-to-locule cavity size showed the most negative correlation in both early (−0.76) and late season (−0.71) measurements. The most important differences in correlation between early and late measurements were the loss of significance in the correlation when comparing compression force with fruit size, total pericarp area, and locule size (mean decrease in correlation of 0.25). Also, the correlation between total pericarp area and outer pericarp diameter changed from 0.11 (non-significant at *p*=0.05) in the early season to 0.38 (significant, *p* < 0.05) in the late season.

### 3.2. Changes during ripening process at cultivar level

Our image-based methodology allowed us to precisely quantify changes in fruit internal structure and firmness during the ripening process. Moreover, fitting quadratic models on the available data allowed us to predict estimates for either intermediate or future time points (Figure 3). Here, we computed predictions for each cultivar, over a 45-days period with data points every day (10 totals).

**Figure 3.**
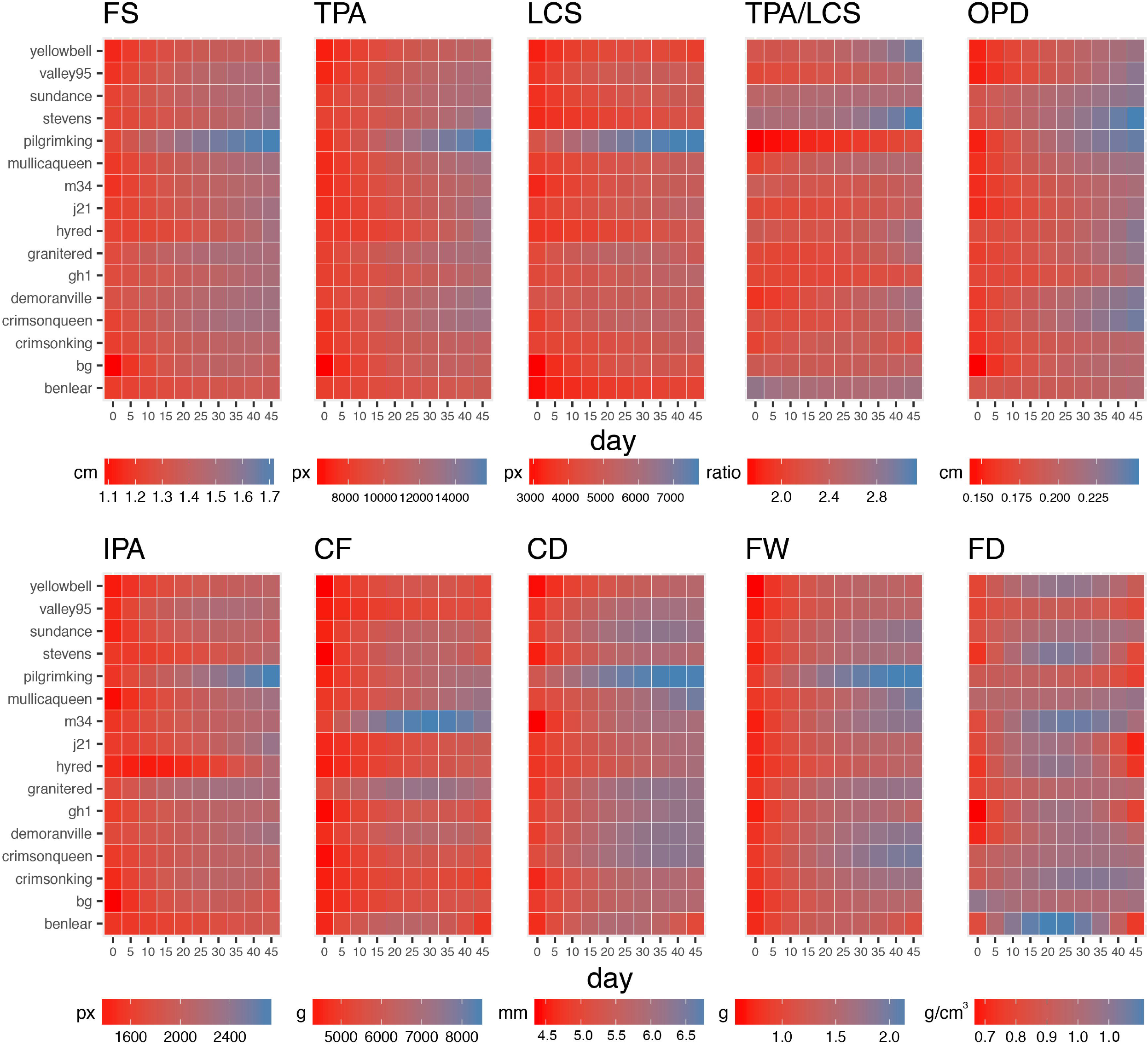
Changes in fruit structure and firmness attributes during the ripening process at cultivar level. Heatmaps for fruit size (FS), total pericarp area (TPA), locule size (LCS), total pericarp area-to-locule size (TPA/LCS), outer pericarp diameter (OPD), internal pericarp area (IPA), compression force (CF), compression distance (CD), fruit weight (FW), and fruit density (FD) are colored according with fitted values obtained from a quadratic model evaluated every 7 days during 10 time points. Coefficients and significance are provided in Supplementary File 2.

Consistent with field observations, at day 45 (last predicted data point), ‘Pilgrim King’ had the largest mean fruit size with 1.72cm, followed by ‘Crimson Queen’, ‘Demoranville’, and J21 with 1.52cm each. Interestingly, ‘Pilgrim King’ reached 1.52cm at day 15, almost a month before the previously mentioned cultivars. ‘Ben Lear’, ‘BG’, and Yellow Bell had the smallest fruit at 45 days with a width less than 1.40cm.

Regarding TPA/LCS ratios at day 45, we observed that ‘Pilgrim King’ (the cultivar with the largest fruit size) had the smallest ratio with 2.11, whereas ‘Stevens’, which had smaller size fruit (1.51cm), had a TPA/LCS ratio of 3.12. ‘Ben Lear’ and ‘Yellow Bell’, which also had small fruit size, had a TPA/LCS ratio of 2.68 and 2.96, respectively. Unexpectedly, some cultivars showed different TPA/LCS tendencies during the ripening process; for example, ‘Demoranville’ had an increase of 41% when comparing the first and last time points, whereas ‘Ben Lear’, ‘M34’ and ‘Sundance’ had little to no change (±5%).

As mentioned before, OPD had a moderate correlation with fruit size (*R*^2^=0.66), indicating that larger fruit tend to have larger OPDs. However, ‘Stevens’ did not conform to this tendency having the largest OPD (0.25cm) even when its fruit size was around the average for that time point (1.51cm, day 45). As reference, the OPD for most of the cultivars accounts for 15% of the total size diameter, whereas for ‘Stevens’ and ‘Pilgrim King’, it was 17% and 14%, respectively. At day 0, ‘Ben Lear’ and ‘Sundance’ showed an OPD comparable with ‘Stevens’ (>0.18mm, ‘Pilgrim King’ had 0.15mm); however, they showed a slow increase later in the season.

Both firmness traits showed contrasting results between each other and with some of the stamp-based descriptors. Unlike other traits, the maximum value for compression force was not observed at later time measurements, but on day 30 (‘M34’, 8472.94g); cultivars like ‘Ben Lear’, ‘GH1’, and ‘Stevens’ showed the same behavior (*beta2* significance *p* < 1×10^−4^). At day 45, ‘M34’ and ‘Mullica Queen’ had CF values greater than 7000g, whereas ‘Ben Lear’ and ‘Crimson King’ had values lower than 5000g. Compression distance showed a gradual increment in Pearson’s correlation with fruit size, from 0.36 at day 0, to 0.91 at day 45.

### 3.3. Grouping cultivars with similar fruit characteristics

To better understand the cultivar trait dynamics across developmental times, and to characterize groups of cultivars that perform similarly, we carried out a principal component analysis (PCA) at each evaluation time point, based on all of the stamp and firmness descriptors combined. In general, for PC1-3 (Figure 4A), we found little separation between cultivars early in the season (up to day ~22), but towards later dates certain trends were observed, and the separation of some cultivars became evident. For PC1, ‘Pilgrim King’ separated considerably well (increased in score) from the rest of the cultivars, whereas ‘Ben Lear’, ‘Yellow Bell’, ‘BG’ and ‘Crimson King’ separated in the opposite direction, although in lower magnitude. PC1 was consistently affected (i.e., loadings, Figure 4B) during the ripening process by TPA and LCS (although this showed a decrease in later dates); CF showed some increase in the second part of the studied dates, whereas IPA showed consistent but low loadings during the whole season. As shown in Figure 4C, PC1 increased its explained variance from 60% at day 0 to 80% at day 45 (Figure 4C). Based on the trait variation registered by PC2, we observed that cultivar ‘M34’ was separated considerably well from the rest of the cultivars, but the maximum separation point was achieved towards the middle of the season. Towards the end of the evaluated season, some cultivars showed an increasing tendency to separate (i.e., ‘Mullica Queen’). As expected by the separation of ‘M34’, CF showed very high loadings, whereas TPA and LCS showed moderate values. PC2 explained ~25% of variance in early season dates, 35% in middle dates, and decreased to 15% towards the end of the season. In general terms, PC3 showed no clear cultivar separation, although some tendencies suggest that major changes could be visible later in the season (i.e. ‘Crimson King’ and ‘Stevens’). PC3 was majorly affected by TPA and LCS (in opposite directions); CF showed highly negative values at early dates, but towards the middle of the seasons it became almost 0. PC3 explained less than 15% of the trait variation during the whole season. Moreover, a PCA using all time points together showed consistency with the analysis previously mentioned, a clear separation of ‘Pilgrim King’ and ‘M34’ mainly because their differences in fruit size and firmness, respectively (Figure 4D).

**Figure 4.**
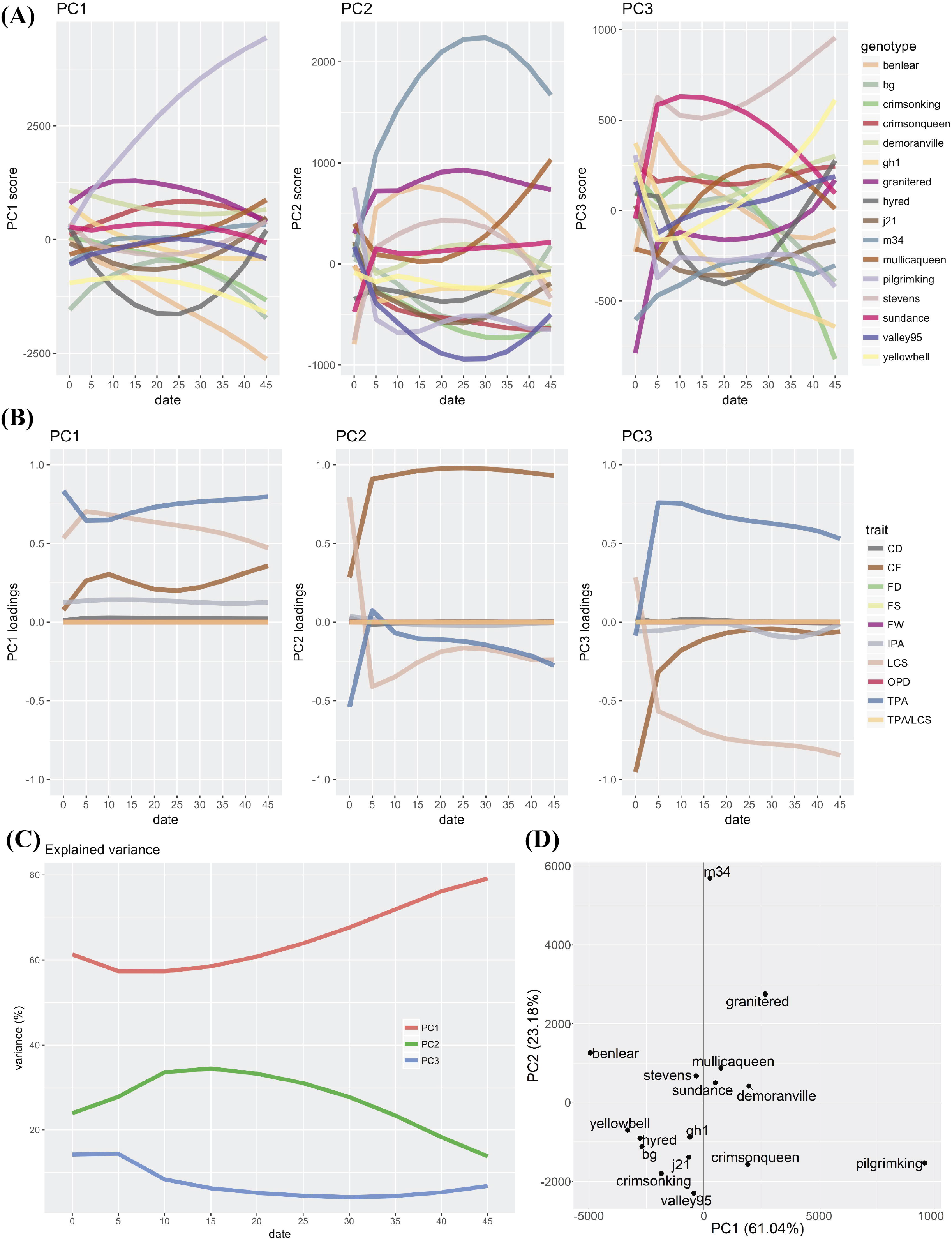
PCA grouping cultivars with similar fruit characteristics. (A) Principal component scores for PC1-3. (B) Loadings for fruit size (FS), total pericarp area (TPA), locule size (LCS), total pericarp area-to-locule size (TPA/LCS), outer pericarp diameter (OPD), internal pericarp area (IPA), compression force (CF), compression distance (CD), fruit weight (FW), and fruit density (FD). (C) Explained variance by component. (D) PCA using all data points at the same time.

### 3.4. Effect of storage on fruit firmness

In this study, we compared the maximum compression force (CF) and maximum compression distance (CD) between fresh fruit and fruit stored at −25°C for 60 days and then thawed at room temperature. In this comparison of fresh and thawed fruit, samples came from fruit harvested on the last two collection dates which were days 33 and 40 (Figure 5). CF of fresh fruit samples harvested at day 33 declined, on average, from 5963.78g to 1483.23g after the freeze-storage period; similarly, for the fresh fruit samples harvested at day 40, CF went from 6381.03g to 1532.01g after the freeze-storage period. CF values for fresh fruit were significantly different between days 33 and 40 (t-test, at *p*<0.05), whereas freeze-stored fruit were not (t-test, at *p*<0.05). For maximum compression distance, the difference between fresh and freeze-stored fruit was subtle for day 33; CD went from 583.75mm to 583.37mm, whereas for day 40, it went from 614.90mm to 582.49mm. As for CF, significant differences (t-test, at *p*<0.05) were only observed when comparing days in fresh fruit. In addition, we calculated the Pearson’s correlation between fresh and freeze-stored fruit for both CF and CD measurements. In both cases, the correlations were significant (at *p*<0.05) with *R^2^* coefficients of 0.77 and 0.90 for CF in days 33 and 40, respectively, and 0.74 and 0.77 for CD in days 33 and 40, respectively. (Figure 5, Supplementary File 2). The high correlation between fresh and freeze-stored fruit firmness measurements reveals that separation among cultivars is maintained after a freeze-storage period (i.e. ‘M34’ showed the highest compression force when fresh and after storage).

**Figure 5.**
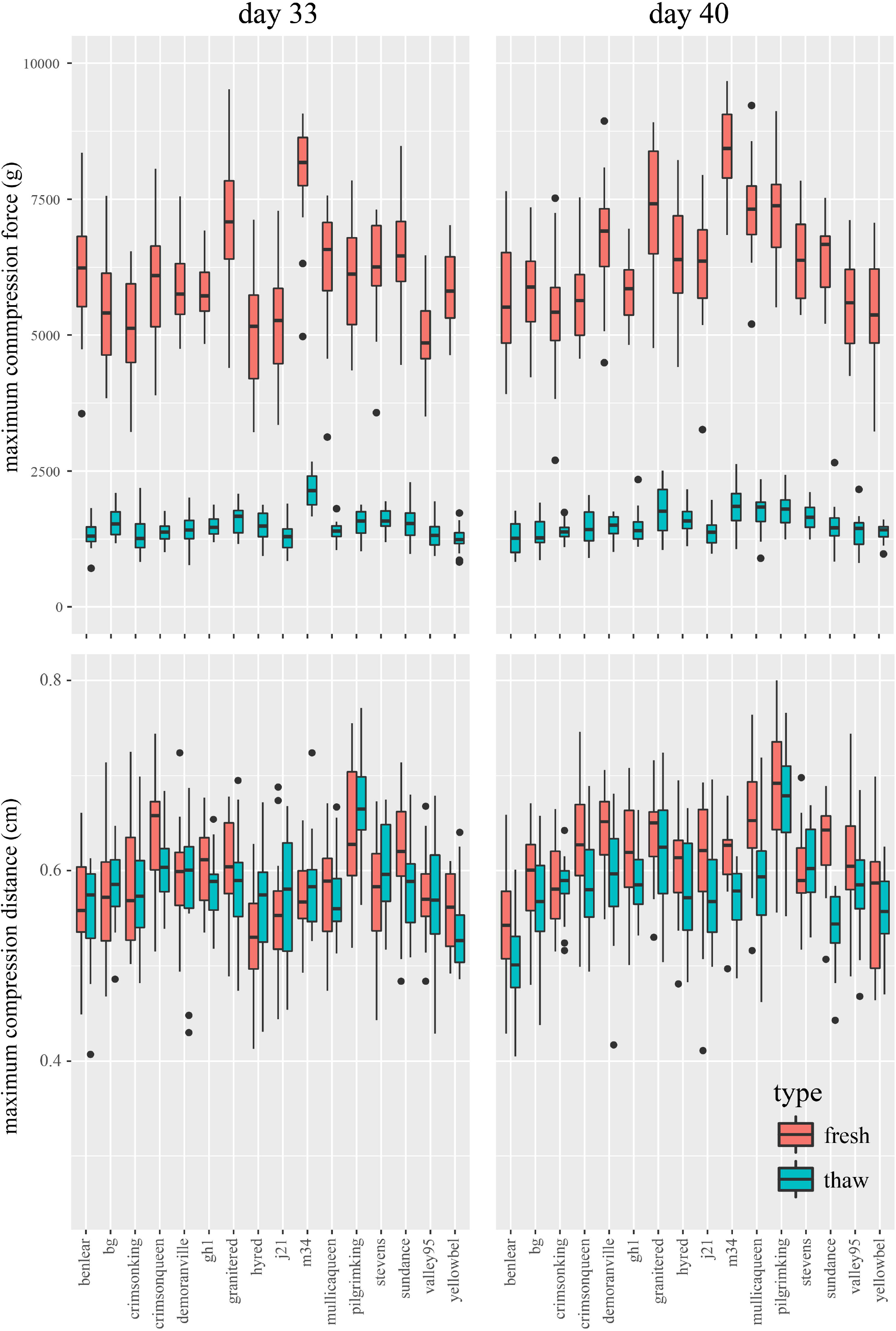
Changes in the firmness properties after storage. Maximum compression force (top panels) and distance (bottom panels) between fresh and stored fruit at two different dates (days 33 and 40).

## 4. Discussion

### 4.1. Measuring internal fruit structure

Most of the studies published in the last decades involving high-throughput phenotyping and computer vision methods have been completely focused on external fruit attributes (Gonzalo et al., 2008; Depypere et al. 2009; Prohens et al., 2012; Zhang et al., 2014; Walter et al., 2015). Fruit size, shape, weight and color are among some of the parameters widely used by researchers, breeders, and the industry. Several phytochemicals, mainly those that confer certain characteristics such as flavor or nutritional value, complete the list of fruit quality traits often considered in cranberry (Diaz-Garcia et al., 2018). However, it is undeniable that fruit firmness and the internal characteristics of fruit play an important role in both industry processing and consumers’ preference. In large fruit crops such as tomato and apple, several studies have been conducted regarding the internal attributes of fruit, their diversity across populations, and how these affect other properties such as storage and firmness (Konopacka and Plocharski, 2004; Brewer et al., 2006). In small crops, little is known about these attributes and the phenotyping technologies for gathering this type of data are lacking.

The methodology for measuring internal structure developed and implemented here is simple to use, inexpensive, and can produce massive amounts of data. Within few minutes, several cranberry samples can be precisely characterized and the data can be directly used either in the industry, research or crop breeding. There are several other alternatives to massively measure the internal structure of fruit. One of the most powerful and non-destructive methods involves the use of computerized tomography (CT) scans, which provides cross-sectional images of any objects interior (Magwaza and Opara, 2014; Jarolmasjed et al., 2016). Although more precise, CT equipment is expensive, difficult to use, and inaccessible for most users. In this sense, our computer-vision strategy presented here represents an alternative not only for the cranberry field, but potentially (with small adjustments) for other fruit crops with similar structures (for example pepper, tomato, papaya, guava, cantaloupe, avocado, etc.)

### 4.2. Implications for the cranberry breeding and industry

Physiological changes in cranberry fruit during the ripening process have been correlated with an increase in fruit size, pericarp area, and locule size, as well as an increase in sugars and phytochemicals such as anthocyanins and flavonols (Vvedenskaya and Vorsa, 2004; Wang et al., 2017) (Çelik et al., 2008). Our results demonstrate that, in addition to the factors mentioned above, fruit internal structure and firmness also changed at different rates across the season for each cultivar tested. Similar to other crops, cranberry cultivars have been developed (or are under current development) to fill specific consumption needs and market niches. As expected, these niches determine the unique set of specifications/requirements that cranberry fruit must meet, and ultimately define the target traits breeders have to select for. For example, ‘Pilgrim King’ had the highest fruit size and weight among all cultivars tested during the entire harvesting period. According to its developer, this cultivar produces extremely large fruit and is high yielding, which could make it attractive for fresh consumption. However, the capabilities of the phenotyping approach developed and implemented in this study allowed us to quantify with high precision that ‘Pilgrim King’ had the lowest total pericarp area-to-locule size ratio, which could affect its use for SDC production. Total pericarp area-to-locule size ratio showed high variability among the cultivars evaluated. This means it must be taken into account for both industry and breeding programs when evaluating cultivars. In the industry, production of SDCs requires fleshy and consistent fruit, with the capability of being infused and that can maintain their structure after being sliced. Because of this, breeding cultivars for the specific purpose of SDC production requires the implementation of phenotyping strategies that allow the gathering of more complex data and from a multi-trait perspective; breeding solely for yield, fruit size or weight could be inefficient in the long-term if the resulting fruit has a low TPA/LCS ratio.

Another tendency observed here that has important implications for growers and industry is that fruit firmness reached its maximum before the fruit stopped ripening. This observation suggests that harvest could be made at early ripening stages, especially when the crop will be used to supply fruit in which firmness is the major interest. Moreover, early harvests could also benefit cranberry fruit transportation. Usually, cranberries are transported and stored in large amounts using large sized containers; therefore, the fruit must be able to handle large compression forces to prevent physiological and/or mechanical deterioration (Forney, 2003). Although the major characteristics currently determining fruit price in cranberry are anthocyanin content and fruit size, our data suggest that the incorporation of additional fruit attributes such a firmness and other structure characteristics into the decision-making process could benefit both the industry and growers.

### 4.3. Limitations of our study

The measurements we obtained in this study were latitudinal representations of the internal fruit structure. Some varieties (e.g., ‘Yellow bell’) show differences in elongation (in the longitudinal axis), and although this characteristic may be mostly determined by genetic factors, it has been observed that environmental conditions as well as different agronomic practices (e.g., fungicide application) can alter fruit shape. Considering this, our fruit size estimates, which assume spherical shaped fruit, were likely underestimated for cultivars presenting certain elongation levels. Additionally, because of the destructive nature of both strategies implemented here (stamp-based phenotyping and firmness analysis), different sets of cranberry fruit were used on each strategy. By doing this, comparisons between fruit firmness and structure attributes can be done using means only, which reduces the number of samples to compare. The same applies to the trait fruit density since the weight measurement used for its computation was obtained from a mean. In the future a non-destructive firmness measurement can be used to test the firmness of an individual fruit and then later that same fruit internal structure can be analyzed through stamping.

There are many texture traits that were not evaluated in this study that are also likely tied to internal fruit structure parameters and may influence cranberry fruit’s overall quality. For example, the elasticity, fracturability, cohesiveness, and toughness of the berry’s flesh and the elasticity, toughness and hardness of the berry’s skin are all additional texture traits that could be measured in cranberries. No set methodology was used in this study to measure fruit firmness because there currently is no standard methodology for testing texture traits in cranberry. The development of a cranberry firmness methodology in the future would help to set standard test parameters that would make comparing different trials easier and set guidelines for obtaining accurate measurements.

### 4.4. Conclusions and further directions

With the implementation of the computer-vision method developed here, we provide a more complete set of fruit quality descriptors that can be used by the cranberry industry and growers. This is particularly important when deciding which cultivars to use for SDC production as well as for better harvest timing. Moreover, expanding the range of phenotypic attributes considered by breeders could benefit the decision-making to develop improved varieties for processing. Without a doubt, high-throughput phenotyping approaches, like the one presented here, will be key for producing massive and reliable data, especially in large-scale scenarios (i.e., industry). Although exploratory, our study highlights the need of characterizing cranberry cultivars using a multi-trait perspective. Further studies regarding fruit structure and firmness must address the implications of cuticle thickness and how this can relate to firmness properties. In addition, including measurements of seed number and weight could provide important insights in how fruit internal structure, overall size, and firmness are correlated.

## Supplementary Data

**Supplementary File 1.** Dataset containing raw data and predicted values.

## Author contributions

LDG, LRB, AA, and JZ design the study; LDG, LRB, MP and ALH performed the experiment; EG provided materials. LDG and LRB analyzed the data; LDG, LRB, AA, MP and JZ wrote the manuscript.

## Acknowledgments

We want to thank Eric Weismann, Walter Salazar and Abigail Kotecki for their tremendous hard work harvesting, processing and stamping thousands of cranberry fruit. JZ would like to express his gratitude through Ps 136:1. We also thank the anonymous reviewers who helped enhance the quality of this paper.

